# Cortical latency predicts reading fluency from late childhood to early adolescence

**DOI:** 10.1101/2025.02.27.640126

**Authors:** Fang Wang, Quynh Trang H. Nguyen, Blair Kaneshiro, Anthony M. Norcia, Bruce D. McCandliss

**Author notes:** Corresponding author: Bruce D. McCandliss,. Graduate School of Education, Stanford, CA 94305 USA. These authors contributed equally to this work (Fang Wang, Quynh Trang H. Nguyen). Conflict of Interest Statement: The authors declare that the research was conducted in the absence of any commercial or financial relationships that could be construed as a potential conflict of interest. Ethics Approval Statement: The study was approved by the Institutional Review Board of Stanford University. As participants were children, a parent or legal guardian of each participant reviewed a written description of the study and gave written informed consent before the session; each participant also assented to taking part in the research.

## Abstract

The development of fluent literacy skills from childhood to adolescence is strongly constrained by the temporal dynamics of word recognition. Capturing the neural basis of these subtle timing changes in word recognition has remained challenging with EEG measures that lack reliability at the individual subject level. Here, we leverage phase information from Steady-State Visual Evoked Potentials (SSVEPs) to derive precise and reliable temporal dynamics of neural signatures underlying visual word form recognition at the individual level and examine their relationship to reading fluency and comprehension. Typically developing readers (N = 68), aged 8–15 years, viewed a stream of four-character stimulus strings presented at 3 Hz. Significant SSVEP signals emerged for nearly all participants. Signals at 3, 6, and 9 Hz harmonics exhibited a phase pattern consistent with a delay model, indicating a mean latency of approximately 170 milliseconds. Individual variations in latencies demonstrated (a) high internal consistency (*R* = .94); (b) stability across variations in letter string forms (familiar words, nonwords with familiar letters, nonwords with unfamiliar pseudo-characters); (c) a linear relationship with age; and most remarkably, (d) a predictive relationship with individual variation in reading fluency and reading comprehension. These results establish SSVEP visual word form latency as a promising approach for investigating the neural basis of reading development, paving the way for future translational applications in education and offering potential solutions to broader societal challenges in promoting population-level reading fluency.

## 1 Introduction

Reading fluency, which extends beyond mere accuracy measures, plays a pivotal role in both reading comprehension and overall literacy development (Yildiz et al., 2014). While various factors influence reading fluency across individuals (Pennington, 2006), word recognition speed stands out as a unique and significant predictor of reading ability (Landerl & Wimmer, 2008), independent of phonological processing and general performance skills (Macmillan & Creelman, 2004). Therefore, understanding the (neuro)temporal dynamics underlying visual word form recognition is crucial for a robust understanding of childhood reading fluency development.

Electroencephalography (EEG) holds promise for capturing the timing of neural dynamics of visual word recognition processes with millisecond-level precision. Attempts to study the peak timing of Event-Related Potential (ERP) components, such as N170, have been shown to be insensitive to developmental differences in reading development (Amora et al., 2022). In contrast, Steady-State Visual Evoked Potential (SSVEP) paradigms—where a sequence of stimuli are presented at a predefined periodic rate (e.g., 3 Hz)—provide robust and high signal-to-noise ratio (SNR) responses in nearly every subject with as few as 3 minutes of stimulation, and have been linked to reading development (van de Walle de Ghelcke et al., 2021; Wang et al., 2022).

The phase of SSVEP signals has proven to be highly informative in studies of the precise timing of neural processes. In 1970, Da Silva and colleagues calculated what has become to be called “apparent latency” of early cortical processes for vision by examining the slope of the *phase-vs-stimulus temporal frequency* function (Da Silva et al., 1970) after having presented the same stimulus condition at multiple temporal frequencies. When the phase-by-frequency plot is linear over a substantial range, the apparent latency can be calculated using the following formula:

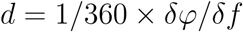

Here, *d* is the apparent latency in seconds, *δφ* is the change in phase over the frequency range in degrees, and *δf* is the change in frequency in Hz. *δφ/δf* is the slope of the phase-versus-frequency plot.

Yeatman & Norcia (Yeatman & Norcia, 2016) extended this phase-by-frequency latency method to study the temporal dynamics of word-selective SSVEP responses. They alternated full-screen images of scrambled and intact text across a range of frequencies from 1-12 Hz. Calculating the slope across these stimulation frequencies revealed early (141–160 msec) and late (>250 msec) left lateralized text-selective responses.

Presenting a single stimulus condition at multiple frequencies, as in Yeatman & Norcia (Yeatman & Norcia, 2016), is both time consuming in general and arduous in particular for children. Fortunately, a more efficient method has recently been validated, taking advantage of SSVEP paradigms that elicit multiple harmonics of the stimulus frequency, thereby providing the necessary frequency range to model latencies using the same procedures. Norcia et al. (Norcia et al., 2020) noted that the phase of the SSVEPs to contrast-modulated luminance probes increased linearly over the first several harmonics of the stimulation frequency. This linear function was then used to derive the latency for the SSVEPs using the *phase-by-harmonic* slope that emerged when results were averaged across the group of participants. The assumption underlying this approach is that the harmonics of the group-level evoked response come from a single cortical source with a fixed delay. The current study was designed to validate and extend this approach by fitting phase-by-harmonic slopes to multiple harmonics at the single-subject level.

Prior SSVEP work has used the “oddball” paradigm—where a sequence of *base* stimuli (e.g., pseudofont strings) are periodically replaced by contrastive *oddballs* (e.g., words) at a slower predefined rate. SSVEP responses emerging at the slower oddball frequencies are thought to reflect categorical discriminations related to higher-level visual processes such as visual word recognition, while the responses at the faster base frequency (and its harmonics) are often interpreted as only reflecting lower-level sensory processes that are insensitive to visual word form processing (Lochy et al., 2016; Volfart et al., 2021). However, recent SSVEP studies using a 3 Hz base stimulation rate across adults and children have revealed multiple neural sources underlie the base response, some of which are consistent with higher-level visual word processing in regions such as the visual word form area (VWFA) (Wang et al., 2021, 2022). Such results highlight the importance of analyzing base responses in greater detail, as this could enhance our understanding of the neural basis of both general visual and word-related processes and their relationship with reading ability, potentially informing future research in this area.

Examination of the temporal dynamics of the base response is also merited by recent converging evidence suggesting that many aspects of developmental challenges in reading are linked to subtle temporal processing deficits (Stein, 2023). For example, McLean et al. found that word recognition speed is a unique and significant predictor of reading ability, independent of phonological processing or domain-general cognitive skills (McLean et al., 2011). Accordingly, the current study examines the temporal dynamics of the base responses using the phase-by-harmonic function validated in (Norcia et al., 2020) to derive cortical latencies of visual word form processes. Critically, this approach goes beyond modeling the central tendencies of group-level SSVEP responses, but also derives individual subject-level responses, which enable a rigorous examination of the connection between these cortical latencies and measures of reading skills in children at multiple level of analysis.

## 2 Methods

### 2.1 Participants

Data from 68 typical developing children (3rd-8th grade, between 8 and 15 years old, median age 11 years, 36 females) were analyzed. Data from three additional children were collected but not analyzed here due to EEG data-quality issues (i.e., excessive movement during recording, incomplete EEG data, N = 2) and having cataract (N = 1). All participants had normal or corrected-to-normal visual acuity and had no diagnosed reading disabilities. After the study, each participant received a small toy for their time.

### 2.2 Behavioral Assessments

Each participant completed a 30-minute behavioral assessment either before or after the EEG session. All children were tested for handedness (Edinburgh Handedness Inventory, Oldfield (1971)), silent reading efficiency (i.e., speed and accuracy) and comprehension (Test of Silent Reading Efficiency and Comprehension, TOSREC, Wagner et al. (2010)), word reading efficiency (Test of Word Reading Efficiency, Second Edition, TOWRE-2, Torgesen et al. (2012)), and non-verbal intelligence and fluid reasoning (the Matrices subtest of the Kaufman Brief Intelligence Test, Second Edition Revised, KBIT-2, Kaufman (1990)).

### 2.3 Stimuli

Three types of stimuli—words, nonwords, and pseudofonts—were used, each comprising four elements (letters or pseudoletters). There were 30 exemplars for each type of stimulus, for 90 exemplars in total.

All words were common monosyllabic singular nouns that started and ended with consonants. They were chosen to have relatively high frequency (averaged 97.7 per million) with limited orthographic neighbors (average 2.3, range from 0 to 4) based on the Children’s Printed Word Database (Masterson et al., 2010).

Nonwords were generated on an item-by-item basis by semi-randomly rearranging the letters of a corresponding Word while ensuring that they also started and ended with a consonant. Each nonword was meant to be unpronounceable and not follow English phonological rules. Nonwords did not differ significantly from the words on average unconstrained bigram frequency (*t*(55) = *−*0.59*, p* = 0.55).

Pseudofonts were the same words stimuli presented in Brussels Artificial Character Set font (BACS-2 Serif font, Vidal & Chetail (2017)), a strictly controlled set of artificial characters that were designed by manipulating the visual strokes of a letter while removing their associated characters.

Using these three types of stimuli, three experimental conditions were investigated. Conditions A and C involved Word embedded in a base stream of pseudofonts (Word & pseudofont) and nonwords (Word & nonword), respectively. For condition B, nonword were embedded in a stream of pseudofont (nonword & pseudofont). The three conditions were completed in a randomly assigned order and were presented at a 3 Hz base stream Figure 1.

**Figure 1:**
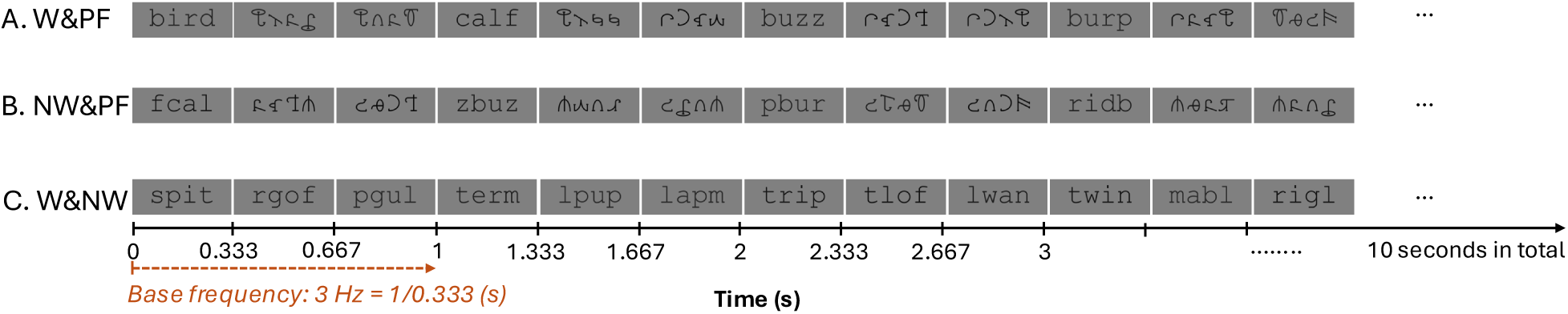
Experimental Design. Example sequences of stimuli presented in the experiment. 3 total presentations per second result in a 3 Hz base stream in all three conditions. A: Word (W) and pseudofont (PF) condition; B: Nonword (NW) and pseudofont (PF) condition; C: Word (W) and nonword (NW) condition. All stimuli were presented centered on the screen.

### 2.4 Experimental Procedure

Prior to the EEG experiment, the procedure was explained to the participant, after which the participant performed a brief practice session.

Participants were seated 1 meter away from the computer monitor in a darkened room during their EEG session. Each condition started with a blank screen shown for a random interval of 1500–2500 ms. During this time, participants were asked to fixate on a white cross in the middle of the screen. Then, each second (aka an epoch) includes the presentation of three stimuli. Twelve successive epochs made up one trial. Ten trials were randomly presented for each condition.

In order to maintain participants’ attention throughout the experiment, a font size change detection task was used. During the recording, the participant continuously fixated on the center of the screen, and pressed a button whenever they detected that the font size of the stimulus got bigger. The font size change detection task was on a “staircase” mode, during which the size of font change became smaller as the accuracy increased, or became bigger when the accuracy decreased. Two size change targets occurred in a non-periodic fashion within a trial. Overall, the entire EEG experiment took around 30 minutes including setup, practice, and breaks between trials.

### 2.5 EEG Recording and Preprocessing

EEG data were collected using 128-sensor HydroCell arrays (MagstimEGI), Electrical Geodesics NetAmp300, and NetStation 5.4.2 software. Data were acquired using Cz as reference at a sampling rate of 500 Hz, and impedances were kept below 50 kΩ. Stimuli were presented using an in-house stimulus presentation software.

Before pre-processing, the data were shifted backwards in time to account for a 60 ms delay introduced by NetAmp300/NetStation software. Recordings were filtered in NetStation using a 0.3-50 Hz bandpass filter that was applied twice to minimize power from frequencies outside the filter range. Data were then imported into the in-house signal processing software for further pre-processing.

During pre-processing, EEG data were re-sampled to 420 Hz to ensure that the frame rate of 60 Hz included an integer number of time samples. Sensors were excluded if more than 15% of samples from the sensor exceeded a 60 *µ*V amplitude threshold. Data from these sensors were replaced by the average value from six of their nearest neighboring sensors.

The continuous EEG data were then re-referenced to average reference (Lehmann & Skrandies, 1980) and segmented into 1-second epochs. Epochs with more than 10% of data samples exceeding the noise threshold of 30 *µ*V or any part of the sample exceeding the blink threshold of 60 *µ*V were excluded from the analysis on a sensor-by-sensor basis. Artifact rejection was performed to remove epochs containing artifacts such as blinks or eye movements. If an epoch exceeded the peak/blink threshold in more than 7 sensors, the entire epoch would be removed in all sensors.

Recursive Least Squares (RLS) filters were then used to filter continuous EEG signals in the time domain (Tang & Norcia, 1995). The filters were tuned to each of the analysis frequencies (i.e., 3Hz, 6Hz, 9Hz) and converted to complex amplitude values by means of Fourier transform. Complex-valued RLS outputs were decomposed into real and imaginary coefficients for input to the spatial filtering computations of Reliable Components Analysis (RCA).

The first and last 1-second epochs of each 12-epoch trial—each comprising three stimulus events—were excluded from further analysis. This was done to reduce the influence of transient responses associated with initial exposure to the stimuli and due to more blinking often occurring at the beginning and end of a trial. Thus, analysis was performed on 10 1-second epochs per trial.

### 2.6 Analysis of EEG Data

#### 2.6.1 Reliable Components Analysis (RCA)

Reliable Components Analysis (RCA (Dmochowski et al., 2012, 2015)) is a matrix decomposition technique that derives a set of physiologically interpretable *reliable components* (RCs) by maximizing trial-to-trial covariance relative to within-trial covariance. Since response phases of SSVEP are constant over repeated stimulations, RCA uses this trial-to-trial reliability to decompose the entire 128-sensor array into RCs, the activations of which reflect phase-locked activities.

Given a sensor-by-feature (i.e., real and imaginary Fourier coefficients) data matrix, RCA computes linear weightings of sensors such that the resulting projected data would exhibit the maximal Pearson Product Moment Correlation Coefficients (Pearson, 1896) across trials. Each RC is a linear combination of sensors that can be visualized as scalp topographies using a forward-model projection of the eigenvectors (spatial filter vectors) (Parra et al., 2005).

Relative to other spatial filtering approaches such as Principal Components Analysis (PCA), RCA achieves high output SNR with a lower trial count (Dmochowski et al., 2015). This is especially useful for investigations involving children since it reduces the amount of data required to observe a robust signal, thereby shortening the EEG data acquisition time.

#### 2.6.2 RCA Calculation

To test brain responses to visual word form stimuli, we computed RCA at the base frequency and its harmonics. Specifically, we input the real and imaginary frequency coefficients of the first three harmonics of the base frequency (i.e, 3 Hz, 6 Hz, and 9 Hz) over the 128-sensor array. To enable a direct quantitative comparison of the three conditions in a shared component space, we computed the RCA weights over the three conditions together.

#### 2.6.3 Analysis of Component-Space Data

To choose which RCA components to examine, we first assessed statistical significance of each component’s eigenvalue coefficient via permutation test (Supplement Figure S1). The permutation test involved generating a null distribution of eigenvalue coefficients against which to compare the observed (intact) RCA coefficients by performing the RCA on 500 versions of surrogate “phase scrambled” sensor-space EEG data. This method disrupts the across-trials covariance of the data and temporal structure of the signal while preserving autocorrelation characteristics and magnitude spectra (Prichard & Theiler, 1994). Here, the phases were randomized in the frequency domain by means of rotation matrices, as described by (Wang et al., 2022). Following this, we further identified the statistical significance at each harmonic for a given significant component via Hotelling’s two-sample t^2^ tests (Victor & Mast, 1991) on distributions of real and imaginary Fourier coefficients on a per-harmonic, per-component basis. Specifically, for each RC, a 1-second epoch of component-space data contained 6 data points (3 harmonics *×* 2 real and imaginary Fourier coefficients). Component-space data were first averaged across 1-second epochs (10 epochs per trial, for 10 trials in total in present study) on a per-participant basis. Statistical analyses were then performed across distributions of participants. Multiple comparisons were corrected using False Discovery Rate (FDR (Benjamini & Yekutieli, 2001)). Twenty-seven comparisons (3 harmonics *×* 3 components *×* 3 conditions) were corrected for base analyses where RCA was computed over the data from three conditions together.

To report SSVEP amplitudes across multiple significant harmonics, we used the square root of the summed squared amplitudes (“root sum square”, *RSS*) across multiple harmonics to combine harmonic response amplitude (Tlumak et al., 2011). We refer to this value as the *RSS* amplitude, in units of *µV* .

#### 2.6.4 Group- and Individual-Level Latencies of RCs

Following the statistical analyses of RCs, we derived group-level latency calculations by fitting a line through the phases at harmonics with significant responses using a linear regression method, where the slope and intercept were obtained with MATLAB’s `polyfit` function (Norcia et al., 2020). The slope of the line represents the response latency.

In order to examine the relationship between brain processing speed and reading abilities, latency was also calculated at the individual subject level. The high SNR provided by SSVEP facilitates this individual-level analysis. We projected each subject’s sensor-space data through the spatial filter vectors derived from group-level RC analyses, resulting in individual projected data. As with the group-level calculations, we fit a linear regression line when responses were statistically significant in at least two harmonics, with the precondition that including the third non-significant harmonic would not substantially change the slope (i.e., latency).

#### 2.6.5 Validation of Individual-Level Latency

To confirm the reliability of our individual latency estimation method and ensure that the estimated latency represents signal rather than noise, we conducted several comparisons. First, we compared latencies at both group and individual levels to check for consistency in their range. Second, we compared individual latencies across the three conditions using paired sample t-tests. Finally, we assessed the split-half reliability to rule out potential confounds from different stimulus sets/conditions in latency calculations. Specifically, we concatenated all 30 trials (epochs/seconds) from the three conditions and randomly split them into two halves, each containing 15 trials—five randomly selected from each condition. We then applied the individual-level latency calculation methods described in the previous section to obtain latencies for each subject in both halves and computed the correlation between them.

#### 2.6.6 Visualization of Component-Space Data

We visualized the RCA data in three ways. First, we present topographic maps for the spatial filters of each component. Second, we present bar plots of amplitudes (*µ*V) across harmonics, with significant responses (according to adjusted *p_F DR_* values of Hotelling’s t^2^ tests of the complex data) indicated with asterisks. Finally, we present phase values (radians) plotted as a function of harmonic, when responses are significant for at least two harmonics, this is accompanied by a line of best fit and slope (latency estimate).

#### 2.6.7 Analysis of brain-behavior relationships

Brain-behavior analyses were then performed to assess individual variations in the relationship between latency and participants’ reading scores and biological age. The reading scores analyzed were the participants’ raw scores of TOWRE and TOSREC, representing reading efficiency and reading comprehension, respectively. Because latency did not differ significantly between conditions, the averaged latency across conditions was used for this brain-behavior correlation. Permutation test results showed RC1 explained the most reliability (Supplement Figure S1), so only latencies of RC1 were included.

Linear correlations were calculated between individual latencies and the TOWRE, TOSREC scores, and age. When significant correlation was found, outliers were identified and removed using a three standard deviation (3*SD*) criterion, where data points falling more than three standard deviations above or below the mean were excluded from further analyses. Correlation analyses were then performed again to assess whether the significant relationship still held after removal of influential data points. As reading skills are highly correlated with age, multiple linear regression analyses, with age as a co-variate, were further performed to access whether latency is genuinely correlated with reading skills, independent of age. Similarly, multiple linear regression analyses with reading scores as covariates were conducted to access whether latency remains correlated with age after controlling for reading scores.

Given the well-established association between TOWRE (single word level reading fluency) and TOSREC (sentence level reading fluency and comprehension), and parallel findings linking latency to reading, we conducted mediation analyses to assess whether TOWRE mediates the relation between latency and TOSREC. A nonparametric bootstrap method with 500 simulations was used to estimate confidence intervals and p-values for the indirect effects. Standardized regression coefficients were calculated and reported to assess effect sizes.

### 2.7 Behavioral Analysis

Behavioral responses for the font size change detection task served as a measure of participants’ attention during EEG recording. We conducted one-way ANOVAs separately for reaction time and accuracy to determine whether participants were highly engaged during the whole experiment.

## 3 Results

### 3.1 Participants were highly engaged during the whole experiment

Participants completed a font size change detection task as a means of sustaining attention during the EEG session. The mean and standard deviation (SD) of d-prime and reaction time across three conditions are summarized in Table 1. Separate one-way ANOVAs indicate significant condition differences in d-prime (*F* (2, 194) = 6.02*, p <* 0.01). Post-hoc t-tests revealed that size-change detection was better in words&nonwords stream compared to the words&pseudofont stream (*t* = 2.26*, p <* 0.05), and better in the words&pseudofont stream compared to the nonwords&pseudofonts stream (*t* = 1.79*, p <* 0.05). There was no significant difference across three conditions in reaction time (*F* (2, 194) = 2.14*, p* = 0.12).

**Table 1:** d-prime and reaction time (RT) of the font size change task performance during EEG sessions for each of three stimulus streams. d-prime was computed based on the z-transformed probabilities of hits and false alarms. Values are mean(*±*SD).

|  | words–pseudofonts | nonwords–pseudofonts | words–nonwords |
| --- | --- | --- | --- |
| **d-prime** | 1.70($\pm 0.38$) | 1.58($\pm 0.42$) | 1.82($\pm 0.36$) |
| **RT(s)** | 0.62($\pm 0.11$) | 0.61($\pm 0.13$) | 0.58($\pm 0.10$) |

### 3.2 Base stimulation elicits reliable spatial topographies and latencies across multiple harmonics

We performed RCA on responses at 3 Hz, 6 Hz, and 9 Hz in order to investigate neural activity related to processing of category non-specific visual word features with the results being summarized in Figure 2 for the first, most reliable component (RC1, for RCs 2 and 3, see Figure S2 in the Supplement). Figure 2A displays topographic visualizations of the spatial filter out of RCA pooled over stimulus conditions. RC1 was maximal over right occipito-temporal area. The bar plots in Figure 2B present amplitudes of component space data at harmonics in bar plots, with statistically significant responses in all three harmonics (all *p_F DR_ <* 0.05, corrected for 27 comparisons) for all three conditions. *RSS* amplitude comparisons across conditions showed that there is no significant difference between conditions (*F* (2, 203) = 0.05, *p* = 0.95). The line plots of Figure 2B display the best-fit line of phase values across three harmonics, accompanied by group-level response latencies, represented and calculated by the slope of the plot. Latencies are comparable across the three conditions (165*±*2.2 ms, 162*±*1.2 ms, and 164*±*2.2 ms, respectively).

**Figure 2:**
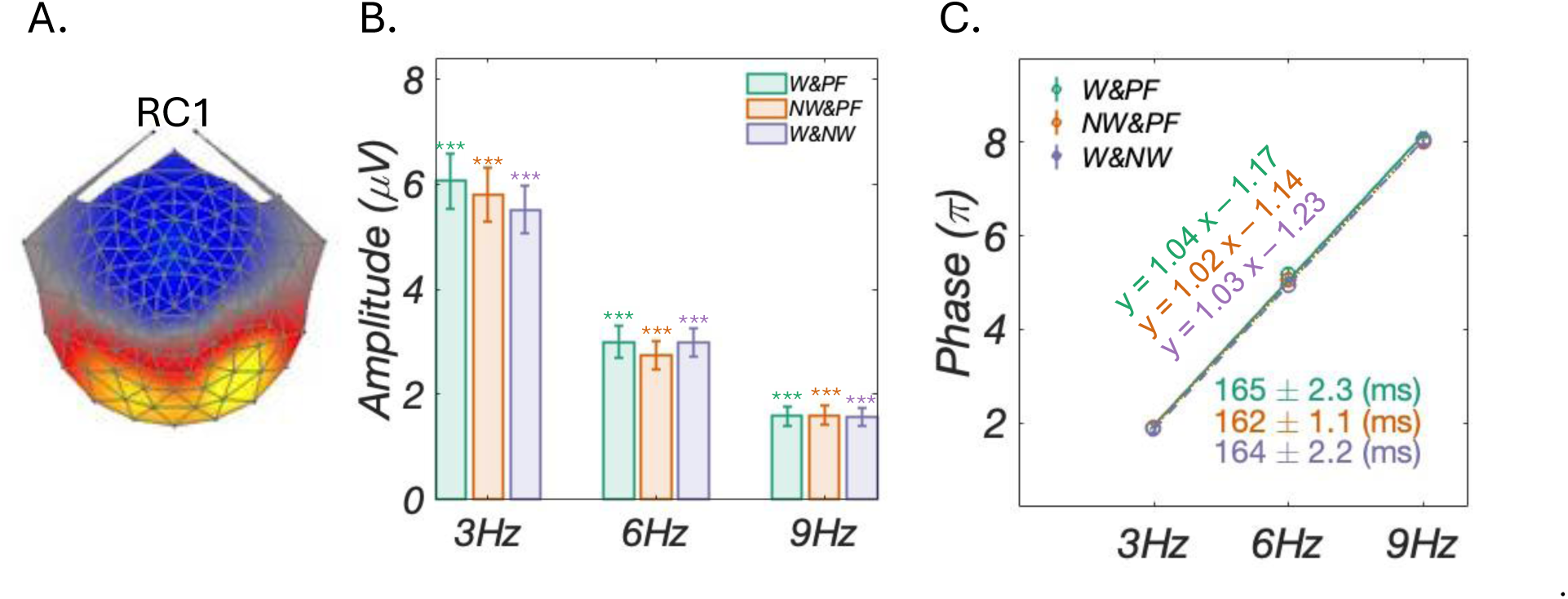
RCA Analyses Results. A: Topographic visualizations of the spatial filter for the first, most reliable components (RC1); B: Amplitude (*µ*V) for each harmonic in bar charts, all response amplitudes are statistically significant (*: *p_F DR_ <* 0.05; **: *p_F DR_ <* 0.01; ***: *p_F DR_ <* 0.001); C: Group-level phase (*π*) and latencies (*ms*) of the three conditions. There are no significant differences across the three conditions (165*±*2.3 ms, 162*±*1.1 ms, and 164*±*2.2 ms, respectively).

### 3.3 Individual-level latencies are highly reliable

At the individual level, latencies of 64 of 68 (94%) participants could be estimated for RC1 (i.e., at least two harmonics had significant signals with the third non-significant harmonic not affecting the slope of line of best fit), allowing for further analyses. The group-level latencies (Supplement Figure S3.A) are broadly similar to the individual-level latencies (Supplement Figure S3.B), both are ranged from 150-250 ms, highly consistent with the well-known 170 ms findings in ERP field. There is no main effect of condition on the individual latencies (Supplement Figure S3.B), and individual-level latencies are significantly correlated between pairs of conditions (all *p <* 0.001 in Supplement Figure S3.C). These findings support the most promising split-half reliability test (*p <* 0.001 in Figure 3D), which demonstrated high reliability of individual latency calculations across different stimulus onsets and various visual word forms.

**Figure 3:**
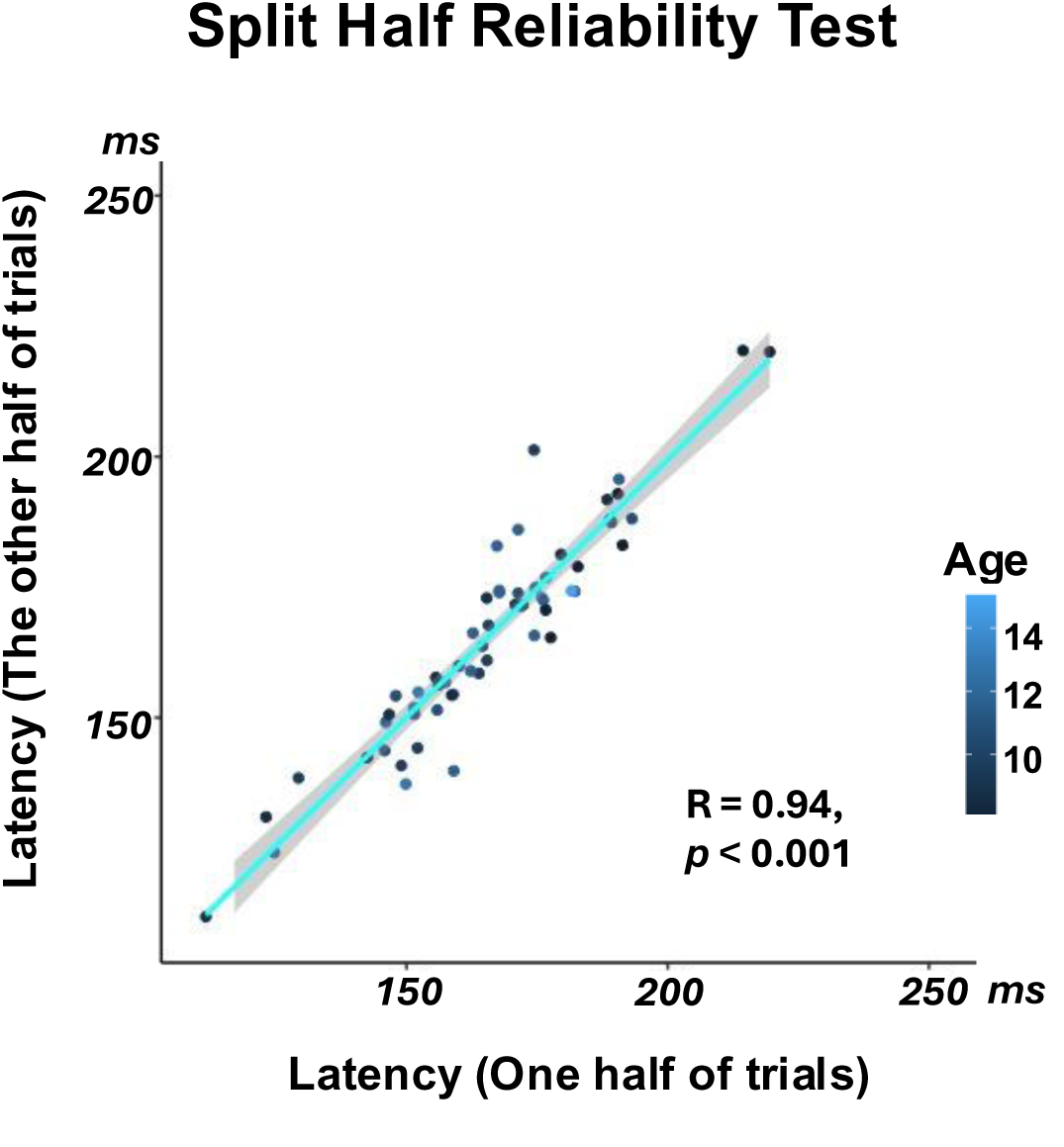
Validation of individual-level latencies. The split-half reliability test was computed to rule out potential confounds from different stimulus sets/conditions in latency calculations. Specifically, we concatenated all 30 trials (epochs/seconds) from the three conditions and randomly split them into two halves, each containing 15 trials—five randomly selected from each condition. The split-half reliability test of all 30 trials across the three conditions demonstrates high reliability, indicating the latencies represent robust signal rather than noise, and more importantly, they are stable across variations in letter string forms.

### 3.4 Individual latencies were negatively correlated with reading scores and age

We tested brain-behavior relationships using the estimated RC1 latencies of individual subjects in conjunction with their reading scores and age. We focus on response latencies only for base processing, as responses at base harmonics were stronger than those at oddball harmonics and most individual-level responses were statistically significant at at least two harmonics (see Methods).

We observed negative correlations for all three comparisons (Figure 4A). Linear correlations between latency and each reading score showed that participants with better word (TOWRE) and sentence (TOSREC) reading fluency and comprehension had shorter latencies. Multiple linear regression correlations showed that the effects of TOWRE (*p <* 0.001) and TOSREC (*p <* 0.01) on latency are still statistically significant when controlling for age. In addition, the effect of age on latency is statistically significant when controlling for reading efficiency and reading comprehension (all *p <* 0.05).

**Figure 4:**
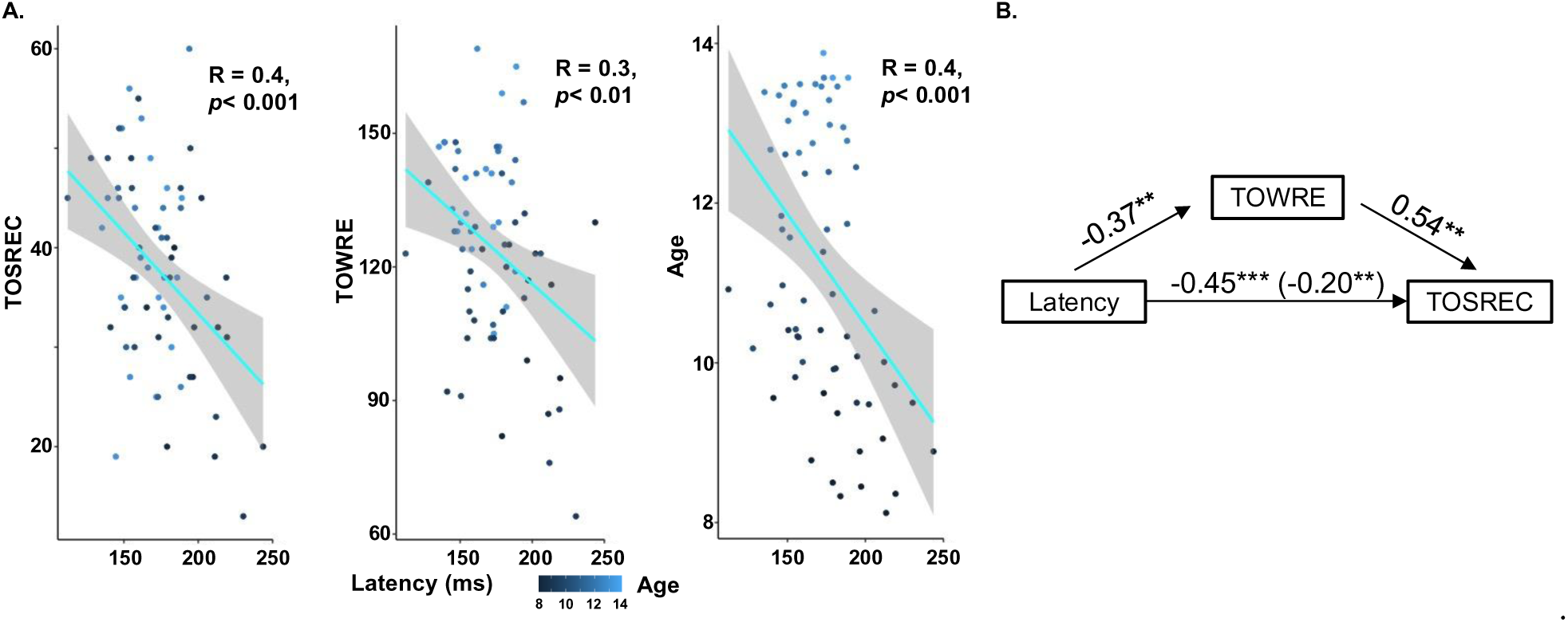
Statistically significant correlations between latency, reading skills, and age, as well as a significant mediation effect. A: Linear regressions between individual-level latency

### 3.5 Relationship between latency and sentence reading fluency is mediated by single word reading fluency

A causal mediation analysis was conducted to examine the effect of latency on sentence reading fluency (TOSREC), with single-word reading fluency (TOWRE) as the mediator. The effect of latency on the TOSREC was significantly mediated via TOWRE. The standardized regression coefficient between latency and TOSREC and the regression coefficient between TOWRE and TOSREC were significant, both *p < .*05 (Figure 4B). The indirect effect was (−0.37) × (0.54) = −0.2. We tested the significance of this indirect effect using bootstrapping procedures (500 samples). The bootstrapped indirect effect was −0.07, 95% CI [−0.15, 0.02] and was significant, *p < .*05.

## 4 Discussion

### 4.1 Latency calculations using SSVEP and RCA

Examining the latencies of spatial components can offer valuable insights into the temporal dynamics of cognitive operations, including processing speed. Unlike commonly used ERP paradigms which typically provide only group-level latencies and require long recording times, this study employed a SSVEP paradigm combined with an RCA approach. With its high signal-to-noise ratio, our SSVEP paradigm requires only a few minutes of stimulation and can provide not only group-level but, importantly, single-subject latencies using an estimation approach that considers phase angles across multiple SSVEP harmonics (Norcia et al., 2020). Additionally, unlike conventional ERP analyses, which usually involve one or a few preselected sensors in a cluster, RCA computes data-driven weighted linear combinations across the entire montage of sensors, thereby enhancing the feasibility of obtaining the individual-level latencies and the discovery of multiple underlying brain sources. Furthermore, by providing latencies, this study extends beyond the majority of previous SSVEP research, which has primarily focused on topographic analysis of response amplitudes.

### 4.2 Group-level latencies and brain sources underlying visual word form processing

Typically, SSVEP researchers use electrode responses at base frequencies as a baseline check, confirming that no significant differences exist between conditions before proceeding to more extensive analyses at the oddball frequency. In contrast with most previous SSVEP studies, including our own prior work that focused on the oddball response (Wang et al., 2021, 2022; Wang, Kaneshiro, et al., 2024; Wang, Toomarian, et al., 2024), here we closely examined the base frequency response. We find that signals at the base harmonics are more robust than those at the oddball frequencies, making them suitable for latency analysis, particularly when extended to the single-subject level. Second, using the data-driven RCA approach, we find multiple brain sources underlie base frequencies, making it promising to analyze these signals in greater detail. This is particularly significant given that the previous literature implicitly assumes the base response, being independent of stimulus category, is likely generated in primary visual areas early in the object-processing hierarchy (Lochy et al., 2016).

RCA of base streams produced three components with differing topographies and time courses (see Figure 2 and Figure S2). Here, we focus on the most robust component, RC1. The finding of brain responses to stimulus streams over occipital (RC1) areas aligns with many previous studies, including those with fMRI (Brem et al., 2009), ERP (Maurer et al., 2006), and SSVEP (van de Walle de Ghelcke et al., 2021) methodologies, which revealed preferential responses to word-like stimuli. The occipito-temporal (OT) sulcus, at a site known as the *visual word form area* that develops with reading expertise, is particularly attuned to written strings (Dehaene & Cohen, 2011). The OT region plays a pivotal role in efficient reading by receiving feedforward signals from early visual cortex and feedback projected from high-level language areas, such as angular gyrus (Lerma-Usabiaga et al., 2018).

Phase-lag quantification of RC1 provides latencies of approximately 160–170 ms. This 160–170 ms latency and the OT activations are consistent with the N170 component, a neuropsychological marker of visual specialization for print, which has been consistently found in previous ERP studies comparing words or word-like stimuli with symbols (Brem et al., 2006; Maurer et al., 2006; Wang & Maurer, 2017).

Therefore, the presence of response components over occipito-temporal regions suggests that the base response is generated in cortical areas that ultimately encode object category. This indicates that object category information could be derived from the base stream, particularly under longer stimulus durations and slower presentation rates (Wang et al., 2022). How these non-selective responses in higher order areas are converted into category specific responses is an important question for future research.

### 4.3 Correlation of individual latencies of visual word form processing with reading and age

The possibility that higher-level orthographic processing could be derived from the base stream opens up opportunities to explore meaningful relationships between these distinctive neural responses and behavioral measures, including reading- and age-related effects.

The higher SNR of SSVEP signals, combined with RCA’s efficiency in extracting phase-locked activities, allows us to obtain highly reliable individual latencies from the majority (94%) of participants, particularly for the most robust component, RC1. This approach enables investigation of individual differences and examination of how latency varies with children’s biological age and reading abilities, providing a new direction to understand cognitive development and literacy skills.

Negative correlations were found between latencies of RC1 and reading skills, as well as age. Children with shorter latency had better single word (TOWRE) and sentence (TOSREC) reading fluency and comprehension, even after controlling for age. Additionally, older children have shorter latencies, even after reading skills were controlled. These results are consistent with several previous findings in the ERP literature of relevance. For example, cross-sectional comparison studies have shown a latency shift from childhood to adulthood due to increased reading skills (Amora et al., 2022; Eberhard-Moscicka et al., 2016; Maurer et al., 2011). Moreover, group comparison studies between typically developing children and those with reading difficulties also provided supporting evidence. For instance, P1 latencies were found to be delayed among dyslexic as compared to normal readers (Breznitz & Meyler, 2003). In addition, abundant research have reported prolonged lower-level visual processing, which may affect further higher-level processing, in readers with dyslexia, particularly in attention demanding situation (Hari & Renvall, 2001) rather than in passive tasks (Regtvoort et al., 2006), presumably reflecting lower quality of automatised performance in readers with reading difficulties.

However, some studies have also found contradictory relationships between latency and reading. In school-aged children, some ERP studies have shown similar mean latencies for both typically developing children and those with reading difficulties or poor readers (Hasko et al., 2013; Kast et al., 2010; Maurer et al., 2011; Zhao et al., 2014). One study even found longer latencies in controls compared with children with reading difficulty (van Setten et al., 2019). This inconsistency also extends to the relationship between latency and age. For example, two studies divided a school-aged sample into young and old subgroups (Maurer et al., 2011; Tong et al., 2016): Maurer et al. (2011) found a longer latency for the younger children compared to the older ones, whereas Tong et al. (2016) reported the opposite pattern. Therefore, further studies are needed to explore the relationship between latency and behavioral measures. However, the consistent patterns across conditions observed in the present study, along with robust split-half reliability, suggest that this approach in the current study may hold promise for future research, particularly when short recording times are required.

### 4.4 Single word reading fluency mediates latency and sentence reading fleuncy and comprehension

We further examined the relationship between latency and sentence reading fluency and comprehension, and found that single word reading fluency mediates this relationship. Our findings provide evidence that latency predicts single word reading efficiency, which, in turn, predicts sentence reading fluency and comprehension. These results align with previous ERP findings showing a latency effect for print, where shorter latencies were observed for familiar single-word recognition in the native language compared to unfamiliar visual scripts from other language systems (Wang & Maurer, 2017, 2020). Increasing processing speed for familiar visual word form processing may reflect increased automatization, driven by reading expertise developed through education and daily exposure (McCandliss et al., 2003), which in turn predicts comprehension and development of text reading. This is an important finding, as it suggests that single-word reading fluency can be explicitly measured and potentially trained through tasks that engage automatic processing, even before formal reading acquisition and instruction begin. Future training studies could lay the groundwork for remediation efforts aimed at improving single-word reading efficiency, ultimately enhancing text reading fluency, comprehension, and overall literacy development.

### 4.5 Challenges and limitations

The individual-level latency estimate approach requires that the signal must meet a specific criteria—the phase-by-harmonic best-fit line must form a linear pattern of phase distribution. This constraint explains why we obtained individual-level latencies only for RC1, the most robust component, and not for other components, including those related to oddball responses.

Additionally, to examine individual differences in cortical latency, a study must include a sufficient number of participants with at least two significant harmonics (provided that including a third non-significant harmonic does not substantially alter the slope) at the individual level for meaningful across-subject comparisons.

The stimulus streams in the current study contained mixed categories of stimuli (e.g., words, pseudofonts) rather than separate streams for each category. However, our split-half reliability test—where all trials across conditions were concatenated and randomly split into two halves—confirmed that this did not pose a problem. Nevertheless, future studies using pure stimulus streams for each category would provide more precise evidence on the neurotemporal dynamics of hierarchical processing and its relationship to reading skills.

## 5 Conclusion

The current study examined latencies of SSVEP base stream processing, a response measure that to date has not received much attention in SSVEP research on object and text processing. Latencies for individual participants were remarkably stable across the three stimulus streams and also highly reliable, as confirmed by split-half reliability testing. Moreover, individual-participant latencies were found to correlate significantly with reading skill and age: Children who are better readers or more developmentally advanced tended to have faster pre-categorical image-level processing speed—i.e., shorter latencies. These findings pave the way for future exploration of the developmental patterns of neural temporal dynamics in children with dyslexia and other reading or learning differences. Early assessment of these patterns could help identify children at risk for developing dyslexia before formal diagnosis, potentially serving as an early diagnostic tool.

## Acknowledgments

We thank the students and their families for participating. We also thank Elizabeth Y. Toomarian, Angie M. Wang, and Lindsey Hasak for their help with data collection.

## 1 Supplementary Material

### 1.1 SI Results

#### 1.1.1 Permutation Test Results

Permutation testing was performed on base RCA coefficients. As shown in Figure S1, RC1–3 coefficients were all statistically significant (permutation test *p_FDR_ <* 0.001, each corrected for 3 comparisons).

**Figure S1:**
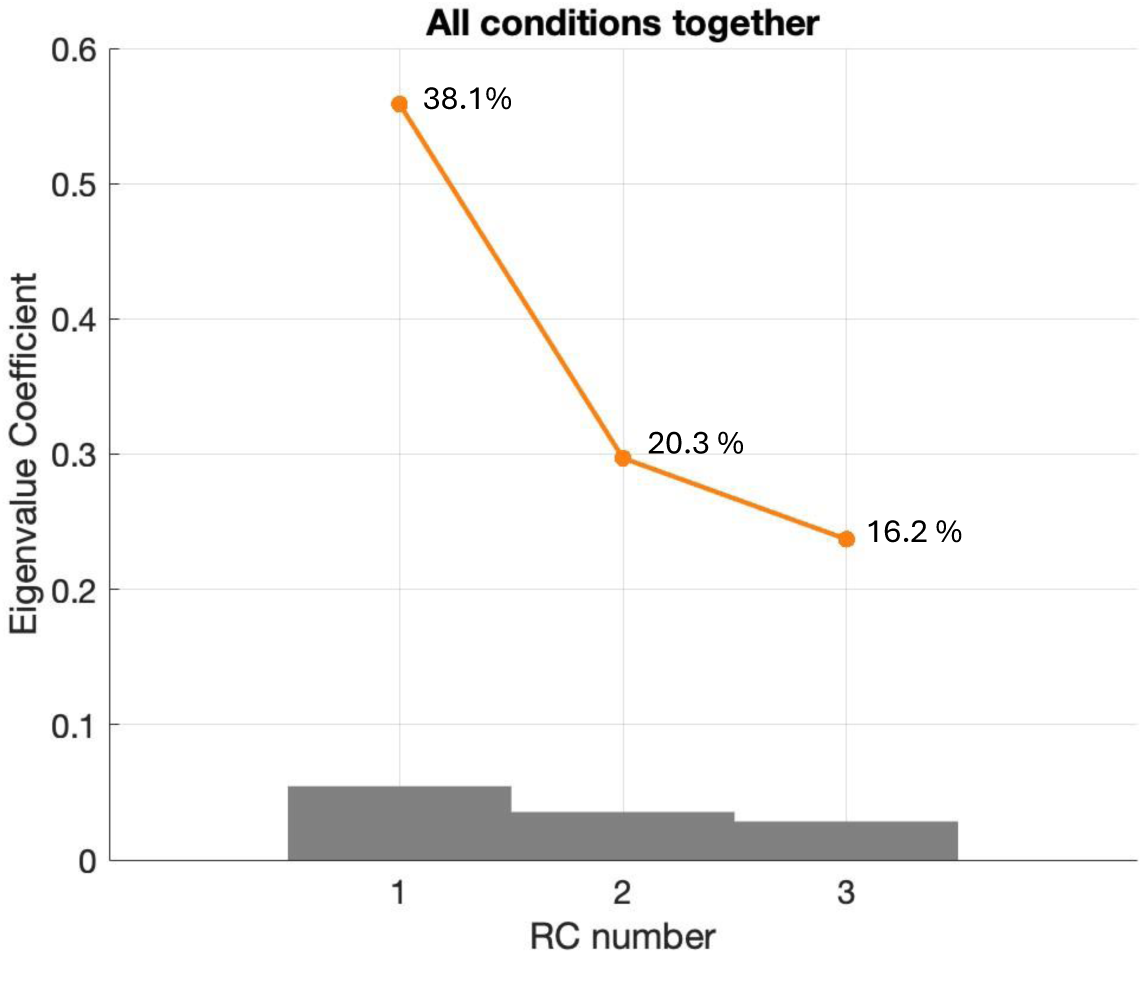
Statistical analyses of eigenvalue coefficients of RC1–3. The line plots denote the distribution of component coefficients for each RCA calculation, and the shaded gray area denotes the 95th percentile of each component’s null distribution. RC1–3 coefficients for base RCA performed on three conditions together were all statistically significant (permutation test *p_F DR_ <* 0.001, each corrected for 3 comparisons). Percentage values represent the percentage of reliability explained by each RC in each condition.

#### 1.1.2 RC2 and RC3 Results

Figure S2A displays topographic visualizations of the spatial filters out of RCA pooled over stimulus conditions for RC2 and RC3. RC2 was displaced to more temporo-parietal electrodes with right lateralization; RC3 showed maximal weightings over medial occipital areas. The bar plots in Figure S2B present amplitudes of component space data at harmonics in bar plots, with statistically significant responses in all three harmonics (all *p_F DR_ <* 0.05, corrected for 27 comparisons) for all three conditions. *RSS* amplitude comparisons across conditions showed that there is no significant difference between conditions (RC2: *F* (2, 203) = 0.10, *p* = 0.90; RC3: *F* (2, 203) = 0.08, *p* = 0.92). The line plots of Figure S2B display the best-fit line of phase values across three harmonics, accompanied by group-level response latencies, represented and calculated by the slope of the plot. Latencies are comparable across the three conditions for each component (RC2: 158–170 ms; RC3: 138–142 ms).

The topography of RC3 lies over early visual cortex and its 140 ms latencies may reflect the dynamics of basic visual feature processing or a very early stage of specialized orthographic processing (Schendan, et al. (1998)). RC3 aligns with the P1 component observed in transient ERP studies, which occurs at around 55-170 ms in children, slightly later than that in adults at around 50-120 ms (Maurer et al. (2005)). This component is also consistent with activation of visual cortex, often the sole activation observed or focused upon in previous SSVEP studies (Lochy et al. (2015); Wang et al. (2021)).

**Figure S2:**
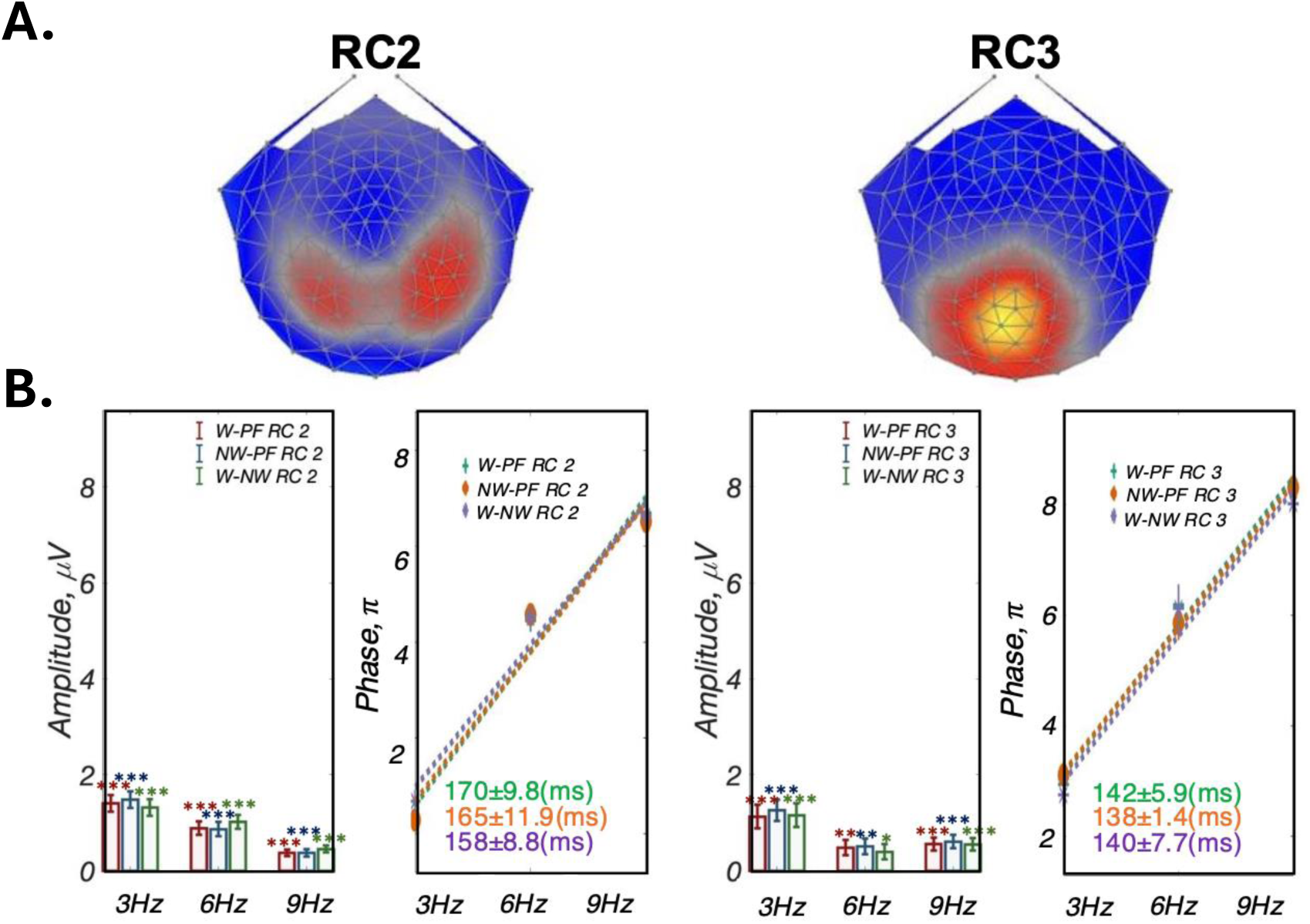
Base RCA Analyses Results for RC2 and RC3. A: Topographic visualizations of the spatial filters (RCs 2&3); B-left: Amplitude for each harmonic at each component in bar charts, all response amplitudes are statistically significant (*: *p_F DR_ <* 0.05; **: *p_F DR_ <* 0.01; ***: *p_F DR_ <* 0.001); B-right: Group-level phase and latencies of the three conditions at each component. There are no significant differences across the three conditions for any of the three components.

**Figure S3:**
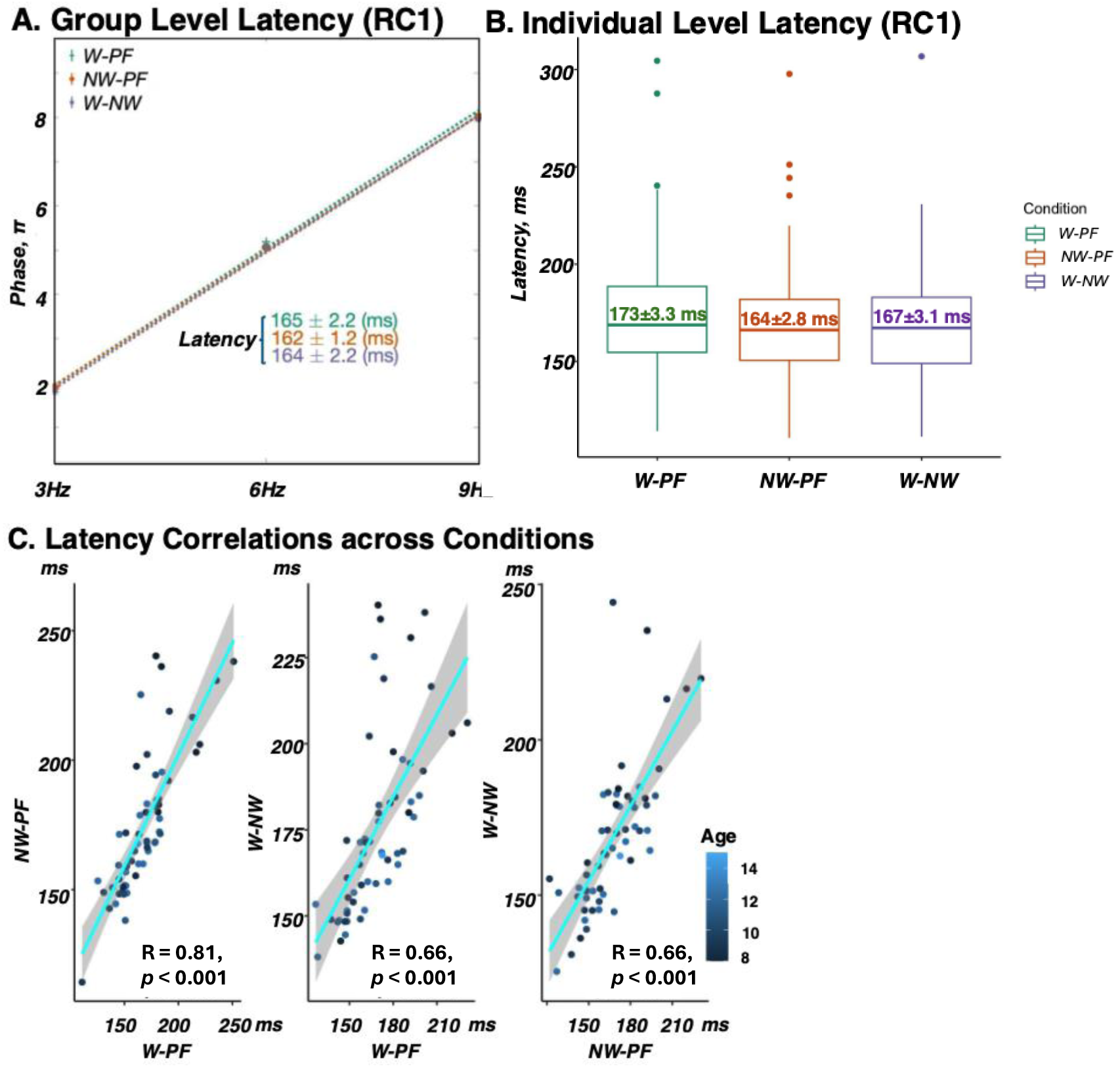
Validation of individual-level latencies. A: Group-level latencies for the three conditions are approximately 165 ms; B: Individual-level latencies across three conditions average between 160-170 ms, slightly different from the group-level latencies but still within a similar range. Individual-level latency comparisons across the three conditions show no significant condition effect (*F* (2, 188) = 1.41, *p* = 0.25); C: Latency correlations across the three conditions reveal high correlations among individual-level latencies within each condition.

### 1.2 Individual-level latencies are reliable

The group-level latencies are approximately 165ms, as shown in Figure S3A. Individual-level latencies, which are broadly similar to the group-level latencies, are shown in Figure S3B: W-PF (*N* = 63; *m* = 173; *SEM* = 3.3); NW-PF (*N* = 64; *m* = 164; *SEM* = 2.8); W-NW (*N* = 64; *m* = 167; *SEM* = 3.1). A one-way repeated-measures ANOVA revealed no main effect of condition on the individual latencies (*F* (2, 188) = 1.41, *p* = 0.25). Moreover, individual-level latencies are significantly correlated between pairs of conditions (all *p <* 0.001 in Figure S3C).

## Footnotes

Abbreviations: Reliable Components Analysis (RCA); steady-state visual evoked potentials (SSVEP).

## Notes

### Competing Interest Statement

The authors have declared no competing interest.

### Summary of Updates

Title was changed to better reflect the whole story of the paper

